# A shared functional organisation underlies vascular disease remodelling

**DOI:** 10.64898/2026.09.03.749124

**Authors:** Alice Bradford, Stefan Bidula, Laszlo Fabian, Derek Warren

## Abstract

Cardiovascular disease involves coordinated remodelling across multiple biological processes. Gene-level signatures vary substantially between studies because different combinations of genes can support similar biological processes. Pathway-level analyses can provide more stable representations of these processes but typically consider pathways independently. We hypothesised that grouping related pathways into conserved biological functions and quantifying their relative weighting would reveal a higher-order, transferable property of vascular tissue that we term functional organisation.

We quantified the relative weighting of six conserved biological functions across independent transcriptomic datasets spanning human vascular disease, experimental models and therapeutic interventions. Vascular tissues exhibited a reproducible functional organisation defined by the balance of these functions. A dominant remodelling trajectory captured coordinated, nonlinear rebalancing of structural, immune, signalling and metabolic programmes, while a second dimension distinguished contractile/ECM organisation from immune activity. This organisation was preserved across independently reconstructed reference cohorts and remained robust to analytical sensitivity testing.

When independent datasets were projected into this fixed framework, biological and clinical phenotypes occupied coherent positions along the remodelling landscape. Plaque-derived vascular, stromal and immune cell populations also occupied ordered functional positions, linking cellular heterogeneity to tissue-level organisation. The same organisational structure generalised across atherosclerosis, peripheral vascular disease and abdominal aortic aneurysm and revealed continuous biological heterogeneity within conventional clinical classifications. Genetic, pharmacological and dietary perturbations reproducibly shifted functional organisation, demonstrating that organisational state is dynamic and biologically responsive.

Together, these findings identify functional organisation as a reproducible tissue-level property of vascular remodelling and provide a framework for understanding cardiovascular disease as coordinated rebalancing of biological functions rather than alteration of individual pathways in isolation.

## Introduction

Cardiovascular diseases remain the leading cause of morbidity and mortality worldwide. Vascular tissues of similar size, composition and pathological classification often exhibit markedly different biological behaviour^1–3^. Across diverse vascular pathologies, including atherosclerosis and aneurysm, disease progression is characterised by coordinated changes in extracellular matrix organisation, inflammatory activation, cellular signalling, metabolism and contractile function. These processes have traditionally been studied through differential gene expression, pathway enrichment, and increasingly through cellular-resolution approaches such as single-cell transcriptomics^4^. Together, these approaches have generated substantial insight into the molecular and cellular mechanisms contributing to cardiovascular disease but do not quantify coordinated tissue-level organisation^4–6^. Consequently, it remains unclear why plaques with similar composition exhibit different stability, why related vascular diseases converge on similar biological behaviour despite arising in distinct vascular beds, and why therapeutic interventions differ in their ability to induce durable remodelling^7^.

Thousands of genes contribute to vascular biology, yet they converge on a relatively small number of coordinated functions required for tissue homeostasis. As vascular tissues simultaneously maintain force transmission, extracellular matrix integrity, metabolism and inflammatory regulation, these biological functions are unlikely to vary independently^8^. If tissue behaviour emerges from the coordinated activity of interacting cell populations, then molecular and cellular measurements should contain higher-order information describing the overall functional organisation of the tissue^9–12^.

This led us to hypothesise that plaque behaviour is an emergent property of the coordinated weighting of multiple conserved biological functions operating across the tissue. We term this tissue-level property functional organisation. Within this framework, vascular states are distinguished by differences in the relative weighting of biological functions. We propose that functional organisation can be quantified by measuring the relative weighting of contractile, extracellular matrix, immune, metabolic, signalling and nucleotide-supply functions within each tissue. These functions represent conserved biological processes that must be coordinated in all multicellular vascular tissues, regardless of disease context. These measurements define a functional signature that provides a quantitative representation of each tissue (Figure 1). Functional signatures can then be projected into a shared plaque remodelling map, from which remodelling position, functional bias, 2D organisational position and reference-state similarity are derived (Figure 1). If present, functional organisation would provide a quantitative description of tissue organisation that complements conventional measures of disease progression and plaque vulnerability. This framework predicts that interventions altering vascular remodelling should induce measurable and directional shifts in functional organisation.

**Figure 1:**
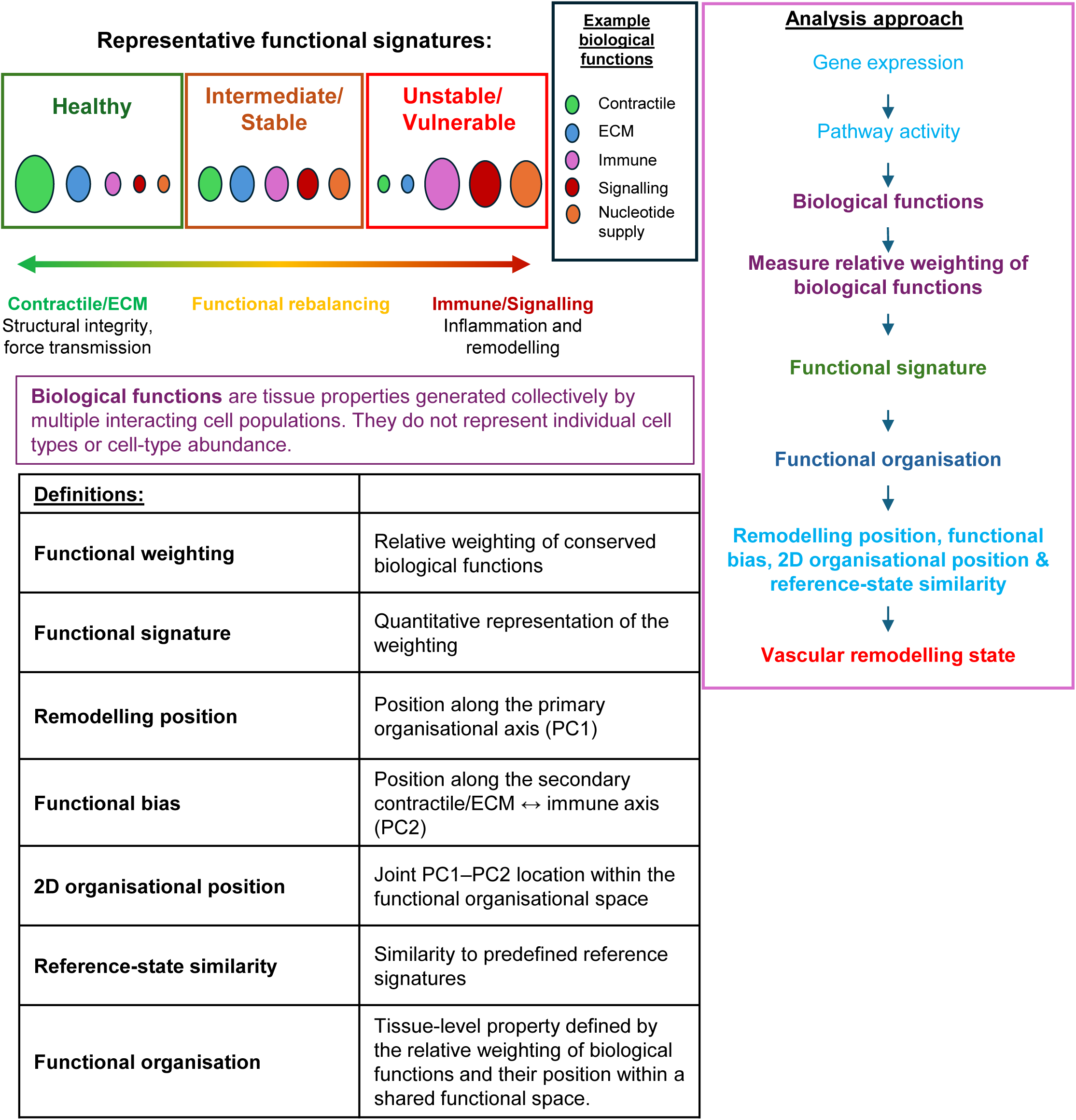
Functional signatures quantify the relative weighting of biological functions. Functional organisation describes the relative balance of biological functions generated collectively across interacting cell populations rather than individual cell types.

In this study, we integrated functional signatures from independent transcriptomic datasets spanning human atherosclerosis, experimental vascular disease and therapeutic intervention. We show that vascular tissues exhibit coherent and biologically interpretable functional organisation that is reproducible across datasets, vascular beds and experimental perturbations. These findings identify a previously unrecognised level of tissue organisation linking molecular mechanisms, cellular phenotypes and vascular remodelling.

## Methods

### Transcriptomic datasets

Thirteen publicly available transcriptomic datasets were obtained from the Gene Expression Omnibus (GEO)^13^. Human bulk transcriptomic datasets representing carotid atherosclerosis were used exclusively to construct the reference plaque remodelling map (Table 1)^14–16^. All remaining datasets, including abdominal aortic aneurysm (AAA), peripheral vascular disease, independent clinical plaque cohorts, plaque-derived single-cell populations and experimental perturbations, were analysed solely as external projection datasets (Table 1)^17–26^. Clinical sample annotations, disease status and intervention groups were retained from the original publications. Thus, independent projection datasets did not contribute to estimation of the reference PCA or its biological-function loadings. This separation between reference construction and external projection was maintained throughout the analysis.

**Table 1:** Datasets used in this study.

| GEO accession | Reference | Tissue/model | Species | Purpose |
| --- | --- | --- | --- | --- |
| **GSE28829** | Döring Y. *et al.* **Circulation** (2012). | Carotid atherosclerotic plaque | Human | Reference plaque remodelling map |
| **GSE43292** | Ayari H, Bricca G. **J. Biosci** (2013). | Carotid atherosclerotic plaque | Human | Reference plaque remodelling map |
| **GSE120521** | Mahmoud *et al.* **Circ Res** (2019). | Carotid atherosclerotic plaque | Human | Reference plaque remodelling map |
| **GSE100927** | Steenman M. *et al.* **Sci Rep** (2018). | Peripheral artery disease | Human | External vascular validation |
| **GSE198600** | Bazan *et al.* **Sci Rep** (2022). | Carotid atherosclerotic plaque | Human | Clinical validation |
| **GSE311535** | Bradford *et al.* **J Mol Cell Cardiol Plus** (2025). | Carotid atherosclerotic plaque | Human | Clinical validation |
| **GSE57691** | Biros E. *et al.* **Oncotarget** (2015). | AAA | Human | AAA validation |
| **GSE98278** | Gäbel *et al.* **J Am Heart Assoc** (2017). | AAA | Human | AAA validation |
| **GSE173974** | Busnelli *et al.* **Heliyon** (2024). | ApoE<sup>-/-</sup> progression | Mouse | Disease progression |
| **GSE56143** | Dunn *et al.* **J Clin Invest.** (2014) | Pharmacological intervention | Mouse | Intervention |
| **GSE234349** | Liu *et al.* **Acta Pharm Sin B.** (2025) | Genetic intervention | Mouse | Intervention |
| **GSE51228** | Maegdefessel *et al.* **Nat Commun.** (2014) | Pharmacological intervention | Mouse | Intervention |
| **GSE159677** | Alsaigh *et al.* **Commun Biol.** (2022) | Calcified atherosclerotic core plaques and patient-matched proximal adjacent (PA) portions of carotid artery | Human | Single-cell projection |

### Generation of functional signatures by single-sample KEGG enrichment

Functional signatures were generated independently for each transcriptomic dataset using a standardised single-sample pathway-analysis pipeline. Each biological sample was analysed individually rather than by case-control differential expression, preserving sample-level variation in functional organisation. Publicly available transcriptomic datasets were downloaded from GEO using GEOquery. Where processed expression matrices were unavailable through the GEO Series Matrix, processed supplementary expression files were identified and reconstructed into sample-by-gene expression matrices. Corresponding phenotype metadata were retained and linked to individual samples. Gene identifiers were converted to Entrez Gene identifiers using org.Hs.eg.db or org.Mm.eg.db and AnnotationDbi, and duplicate identifiers were collapsed to a single expression measurement per Entrez Gene identifier.

For each sample, genes were ranked by expression level, and KEGG Gene Set Enrichment Analysis (GSEA) was performed using clusterProfiler, with pathway sizes restricted to 5–500 genes^27–30^. For the primary bulk-tissue analysis, pathways were retained using a permissive BH-adjusted P threshold of ≤0.5. This threshold was used as a pathway-retention criterion rather than as a test of pathway-level statistical significance and was subsequently examined by sensitivity analysis. A minimum of three retained pathways was required for a biological-function score to be estimable.

KEGG pathways were mapped using predefined keyword dictionaries to six higher-order biological functions representing conserved components of vascular remodelling: Contractile, extracellular matrix (ECM), Immune, Metabolic, Signalling and Nucleotide supply. Because individual KEGG pathways can represent more than one biological function, pathway overlap was explicitly accounted for during module weighting. A pathway assigned to (k) biological functions contributed a weight of (1/k) to each corresponding function. Each biological-function score was calculated as the overlap-adjusted weighted mean of the normalised enrichment scores (NES) of its constituent pathways. This prevented multiply assigned pathways from contributing their full weight independently to several biological functions. The resulting six module-weighting scores constituted the functional signature of each sample.

Where a biological function could not be estimated because of insufficient pathway information, its value was retained as missing (NA) rather than assigned a biological value of zero. Biological missingness was preserved in all stored functional-signature tables. Only for PCA fitting and projection was a temporary computational matrix generated in which missing within-dataset standardised values were replaced by zero, corresponding to the within-dataset standardised mean. This substitution was restricted to the PCA calculation/projection step and did not alter the underlying biological-function scores.

Unlike conventional differential-expression analyses, this approach generated an independent functional signature for each biological sample, allowing healthy/control tissues, disease states, clinical samples and experimental perturbations to be represented within a common functional framework.

### Pathway-retention and normalisation sensitivity analyses

Sensitivity analyses tested whether the reference organisation depended on analytical choices made during construction of the biological-function scores. Pathway-retention sensitivity was assessed by reconstructing biological-function scores across BH-adjusted P-value thresholds of 0.05, 0.10, 0.25, 0.50, 0.75 and 1.00 while retaining the minimum pathway requirement. For each threshold, biological-function scores were standardised within dataset and the reference PCA independently refitted. Solutions were compared with the primary BH-adjusted P ≤0.50 solution after accounting for the arbitrary orientation of PCA axes. Stability was quantified from correspondence of sample coordinates and component loadings and preservation of the dominant organisational structure.

Normalisation sensitivity was assessed by independently reconstructing the reference map following within-dataset z-score standardisation (primary analysis), median–IQR scaling, median–MAD scaling, mean centring without scaling, or no within-dataset normalisation. Each PCA was independently refitted and compared with the primary reference solution after axis alignment.

### Construction and orientation of the reference plaque remodelling map

The reference plaque remodelling map was constructed exclusively from the human bulk atherosclerosis datasets GSE28829, GSE43292 and GSE120521. Biological-function scores were standardised independently within each dataset by z-score transformation before datasets were combined, reducing study-, platform- and scale-specific differences while retaining relative functional variation within each study. PCA was then performed on the six biological-function scores. PCA was fitted once to these reference samples; the resulting loading matrix defined the fixed reference space used for all subsequent analyses. Independent datasets never contributed to refitting the reference PCA. This is a fundamental rule of the analysis pipeline.

The signs of PCA components were fixed after reference construction because PCA orientation is mathematically arbitrary. PC1 was oriented so that increasing values represented movement from healthy towards disease-associated reference organisation. PC1 therefore defined the dominant remodelling position. PC2 was oriented using the biological-function loadings of the fixed reference PCA alone, such that the Immune loading was greater than the mean loading of the Contractile and ECM functions. Consequently, increasing PC2 represents increasing immune weighting relative to contractile/ECM weighting. PC2 was interpreted as a relative functional-balance axis rather than assigning biological meaning solely according to whether individual scores were positive or negative.

The joint PC1-PC2 coordinates defined the two-dimensional functional organisational space. A principal curve was additionally fitted through the reference organisation to characterise nonlinear progression through this space and provide a complementary description of the remodelling trajectory^31^. For analyses requiring direct comparison among independently reconstructed PCA solutions, rescaled PC1 was used as the dominant remodelling coordinate, avoiding instability introduced by independently refitting principal curves.

### Projection of independent datasets

Independent datasets were projected into the fixed reference space without recalculation of the reference PCA loadings. Biological-function scores were first standardised within their source dataset or experimental cohort using the same framework as the reference data and then multiplied by the fixed reference loading matrix. Thus, projection quantified the relative organisation of samples within each independent cohort with respect to a reference coordinate system derived exclusively from human atherosclerosis. Projection datasets could therefore be positioned within the reference organisation but could not alter its axes.

For PCA projection only, missing standardised biological-function values were replaced by zero, corresponding to the mean of the standardised source cohort. The original biological-function values remained NA, and computational substitutions were recorded separately.

### Quantification of functional organisation

Functional organisation was described using complementary quantities that distinguish biological composition from position within the reference map.

#### Functional signature

The functional signature comprises the relative values of the six overlap-adjusted biological-function scores and describes the balance of Contractile, ECM, Immune, Metabolic, Signalling and Nucleotide-supply functions within a sample or population.

#### Remodelling position

Remodelling position is the coordinate along PC1 of the fixed human atherosclerosis reference PCA. PC1 represents the dominant direction of coordinated functional reweighting and is oriented from healthy towards disease-associated organisation.

#### Functional bias

Functional bias is the coordinate along PC2. PC2 represents variation orthogonal to the dominant remodelling axis and is oriented so that increasing values indicate increasing Immune weighting relative to mean Contractile/ECM weighting.

#### Two-dimensional organisational position

Joint PC1-PC2 coordinates represent the location of a sample within the fixed two-dimensional functional organisation and retain both dominant remodelling position and orthogonal functional bias.

#### Reference-state similarity

Early, Intermediate and Late reference organisations were defined from the fixed human reference. Functional signatures from projected samples were compared with the corresponding reference-state signatures using Pearson correlation and cosine similarity, which were equally weighted and transformed by softmax normalisation (temperature, *T* = 0.20) to generate relative signature-match probabilities. Full mathematical definitions are provided in the Supplementary Methods.

### Robustness of the reference functional organisation

Robustness analyses were designed to test distinct potential sources of dependence in the reference organisation: dependence on individual biological functions, their precise relative weighting, individual reference datasets, sample composition and disease-state labels.

#### Functional-definition robustness

Each of the six biological functions was sequentially omitted and the reference PCA independently reconstructed from the remaining five functions. Because omission of a function can cause the first two PCA components to exchange order or rotate within the same two-dimensional subspace, correspondence with the complete reference was assessed both by the best-matching PCA axis and by two-dimensional Procrustes alignment of PC1-PC2 coordinates. This distinguished preservation of the underlying organisational geometry from preservation of an arbitrarily numbered PCA component. The robustness workflow explicitly includes PC1/PC2 axis-swap diagnostics and two-dimensional Procrustes preservation metrics.

Dependence on the precise relative weighting of the six biological functions was tested by 500 random perturbations of module weights. Multiplicative weights were drawn from a log-normal distribution centred on one (SD on the log scale = 0.20) and rescaled within each iteration to a mean of one, thereby perturbing the relative contribution of functions without changing the overall magnitude of the functional matrix. The PCA was reconstructed after each perturbation and correspondence with the unperturbed reference quantified from sample-coordinate correlations, best-axis correspondence and two-dimensional Procrustes correlation.

This analysis tests whether the organisation depends critically on a single higher-order functional category or on exact equal weighting of those categories. It does not constitute random reassignment of individual KEGG pathways among the biological-function dictionaries; such a test would require perturbation before module aggregation.

#### Reference-dataset robustness

Each constituent reference dataset was sequentially omitted, the PCA independently reconstructed from the remaining reference datasets, and the resulting organisation compared with the complete reference. Correspondence was assessed using best-axis correlation and two-dimensional Procrustes correlation, allowing preservation of the organisational subspace to be distinguished from component swapping or rotation. Held-out samples could subsequently be projected into a reference space to which they had not contributed.

#### Sample-composition robustness

Individual-sample sensitivity was assessed using 500 repeated subsampling iterations in which 80% of samples were retained within each study-by-state stratum before independent reconstruction of the PCA. Empirical distributions of correspondence metrics quantified sensitivity of the reference organisation to individual sample composition.

#### State-label permutation testing

To determine whether the relationship between functional organisation and clinical state exceeded that expected by chance, sample-state labels were randomly permuted 1,000 times while the measured functional signatures and reference geometry were retained. For each permutation, linear models quantified the proportion of variance *R*^2^ in PC1, PC2 and principal-curve remodelling trajectory associated with the permuted state labels. The observed *R*^2^ were compared with the corresponding null distributions. Empirical permutation *P* values were calculated as *(k+1)/(N+1),* where *k* was the number of permuted *R*^2^ values equal to or greater than the observed *R*^2^ and *N* was the number of permutations.

#### Diffusion-map embedding

As an alternative nonlinear representation of the functional organisation, diffusion-map embedding was applied to the within-dataset standardised six-function profiles^32^. Principal-component analysis was first performed on the six functional scores without additional centring or scaling, because the input values had already been standardised within their source datasets. The first five principal components, or all available components when fewer than five were available, were used as input to the diffusion analysis.

Pairwise squared Euclidean distances were calculated between samples and converted to a Gaussian affinity kernel, 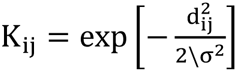. The kernel bandwidth *σ*was estimated as the square root of the median non-zero pairwise squared distance. Self-affinities were set to zero and the resulting kernel was row-normalised to generate a Markov transition matrix. Diffusion coordinates were obtained from the eigendecomposition of this matrix using diffusion time *t* = 1; the first trivial component was excluded and the first two non-trivial components (DC1 and DC2) were retained for visualisation.

Because the sign of an eigenvector is arbitrary, DC1 was oriented to agree with the direction of PC1 in the primary reference analysis. A principal curve was fitted through the resulting two-dimensional diffusion space for visualisation of the nonlinear organisational trajectory. As an additional sensitivity analysis, diffusion embedding was also performed directly on the six-function space without PCA preprocessing, and concordance of the resulting dominant diffusion coordinate with reference PC1 and with the PCA-preprocessed diffusion embedding was assessed.

### Single-cell analysis

Plaque-derived single-cell populations were analysed using differential-expression-derived ranked gene lists rather than single-sample expression profiles. Differentially expressed genes for each cell population were ranked by average log2 fold change and analysed by KEGG GSEA. In this analysis, all returned KEGG pathways were retained for module weighting rather than applying a pathway-level adjusted-P threshold. Pathways were assigned to the same six biological-function dictionaries and overlap-adjusted in the same manner as the bulk analysis. The resulting cell-population functional signatures were standardised across the projected cell-population cohort and projected into the fixed human atherosclerosis reference PCA without refitting its loadings. The scRNA projection script explicitly requires this fixed-reference approach and retains biological missingness as NA.

### Intervention analyses

Independent genetic, pharmacological and dietary datasets were processed independently of the reference datasets. Biological-function scores were standardised within each intervention source dataset and projected into the fixed human reference PCA; intervention samples were excluded from reference PCA training. Changes induced by experimental perturbation were evaluated from changes in functional signature, remodelling position, functional bias, two-dimensional organisational position and reference-state similarity relative to the corresponding experimental controls.

### Statistical analyses

All analyses were performed in R (version 4.6.1) using RStudio (2025.09.02, Build 418). Principal component analysis (PCA) was performed using prcomp() without additional centring or scaling because biological-function scores had been standardised within their source datasets before construction of the analysis matrix. PC1 defined remodelling position and was oriented such that increasing values represented progression from healthy towards disease-associated functional organisation. PC2 defined functional bias and was oriented from the fixed-reference loadings such that increasing values represented greater immune weighting relative to contractile/ECM weighting. The joint PC1-PC2 coordinates defined two-dimensional organisational position. The reference PCA was fitted once to the human atherosclerosis reference datasets and its loadings were fixed for projection of all independent datasets.

A principal curve was fitted through the reference samples to characterise the nonlinear remodelling trajectory and calculate principal-curve pseudotime. Generalised additive models were used to model changes in individual biological-function scores along pseudotime, allowing nonlinear functional reweighting across the remodelling trajectory to be visualised^27,33^. For robustness analyses involving independently refitted PCA solutions, the dominant remodelling coordinate was additionally represented by rescaled PC1 (0–1), enabling direct comparison of trajectory ordering without requiring identical principal-curve geometry between reconstructions.

Agreement between independently reconstructed PCA solutions was assessed using Pearson correlations of component loadings and sample coordinates. Because omission or perturbation of biological functions can rotate the PC1-PC2 space or exchange component order while preserving its underlying geometry, functional-definition and dataset-omission analyses additionally used best-axis correlation and two-dimensional Procrustes correlation. Spearman correlation was used where preservation of ordered reference states was assessed.

Robustness of the reference organisation was evaluated at multiple levels. Study dependence was assessed by sequential leave-one-reference-dataset-out reconstruction. Sensitivity to individual sample composition was assessed using 500 repeated subsampling iterations in which 80% of samples were retained within each study-by-state stratum. Dependence on functional definition was assessed by sequential omission of each biological function and by 500 random perturbations of relative biological-function weights. For weight perturbation, multiplicative weights were drawn from a log-normal distribution with log-scale SD 0.20 and rescaled to a mean of one within each iteration, thereby altering relative functional weighting without changing overall matrix magnitude.

To determine whether associations between functional organisation and clinical state exceeded those expected by chance, sample-state labels were randomly permuted 1,000 times while retaining the measured functional data. Linear models were used to quantify the proportion of variance (R²) in remodelling position (PC1), functional bias (PC2) and remodelling trajectory associated with sample state, and observed statistics were compared with their corresponding permutation-derived null distributions. Empirical permutation P values were calculated as (k + 1)/(N + 1), where k represents the number of permuted statistics equal to or greater than the observed statistic and N the number of permutations.

A two-sided P < 0.05 was considered statistically significant for conventional hypothesis tests. Formal mathematical definitions of functional signature, remodelling position, functional bias, two-dimensional organisational position, reference-state similarity and functional organisation are provided in the Supplementary Methods.

## Results

### A shared functional organisation underlies human atherosclerotic remodelling

Functional organisation was quantified across independent human atherosclerotic transcriptomic datasets encompassing control, plaque, stable and unstable clinical groups. Relative weighting of contractile, extracellular matrix (ECM), immune, metabolic, signalling and nucleotide-supply functions revealed a coherent, low-dimensional organisation of human vascular remodelling (Figure 2A and Supplementary Figure 1). The dominant organisational axis captured coordinated reweighting of these biological functions, while a secondary axis contrasted contractile/ECM organisation with immune activity (Figure 2B and C). Modelling individual functions along the dominant remodelling trajectory further revealed coordinated but nonlinear changes in structural, immune, metabolic and signalling programmes (Figure 2D).

**Figure 2.**
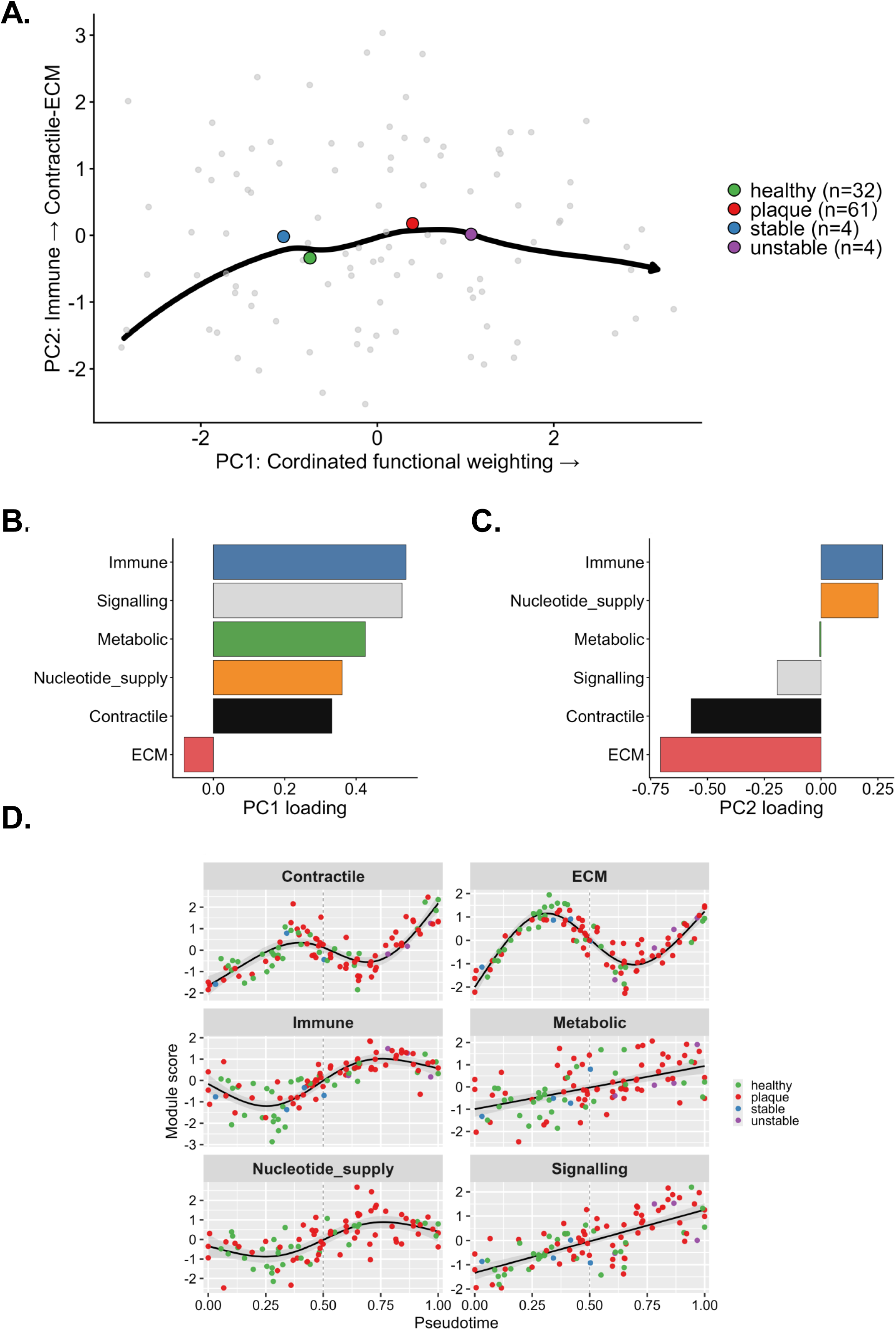
Human plaques occupy a shared functional organisational continuum. **A)** Principal component analysis (PCA) of within-dataset standardised functional signatures. Biological function scores were z-score standardised within each dataset prior to PCA. Each point represents an individual sample, coloured centroids indicate group means for healthy (green, n=32), plaque (red, n=61), stable (blue, n=4), and unstable (purple, n=4) states. A principal-curve fit (black line with arrow) traces the dominant trajectory through the data. **B)** PC1 (x-axis) represents the dominant organisational axis and reflects coordinated reweighting of structural, immune, signalling and metabolic functions. **C)** PC2 (y-axis) captures a secondary functional-balance axis contrasting Contractile/ECM organisation with Immune weighting. **D)** Generalised additive model (GAM) fits of module scores across principal-curve pseudotime show coordinated, non-linear trajectories for each biological function. Each point represents an individual sample coloured by sample state (healthy, plaque, stable, unstable). Lines indicate fitted GAM trends.

We next tested whether this organisation was dependent on analytical choices, individual biological functions or constituent datasets. The dominant organisational axis was essentially unchanged across a broad range of BH-adjusted pathway-retention thresholds and was strongly preserved using alternative within-dataset normalisation approaches (Supplementary Figure 2). Sequential omission of individual biological functions retained the overall organisational structure, and random perturbation of their relative weighting produced maps closely corresponding to the unperturbed reference (Supplementary Figure 3A). Permutation of clinical state labels showed that the observed association of state with remodelling position and trajectory substantially exceeded that expected by chance, whereas functional bias showed a weaker state association (Supplementary Figure 3B). Sequential omission of individual reference datasets similarly preserved the dominant organisation (Supplementary Figure 3C). Diffusion-map embedding recovered the same underlying trajectory using both PCA-pre-processed and direct six-function representations (Supplementary Figure 4).

We fixed this reference organisation and projected all subsequent datasets into it without recalculating the reference loadings. Independent peripheral artery transcriptomic datasets occupied the same functional space across multiple vascular territories, supporting conservation of the organisational framework beyond the atherosclerotic datasets from which it was derived (Supplementary Figure 5).

#### Multicellular plaque populations share a common functional organisation

To determine whether tissue-level functional organisation was reflected across constituent plaque cell populations, we projected independently derived plaque core and proximal single-cell populations into the shared functional organisational framework. Vascular, stromal and immune populations exhibited distinct functional signatures but occupied coherent positions within the same organisational space (Supplementary Figure 6). Fibroblast and endothelial populations showed later remodelling positions, whereas neuronal and VSMC populations were positioned towards the opposite end of the primary organisational axis. Functional bias provided an additional distinction between populations, separating those with greater contractile/ECM weighting from those with greater immune weighting. Reference-state similarity further resolved these differences by quantifying correspondence with Early, Intermediate and Late tissue-level signatures (Supplementary Figure 6). Thus, diverse plaque cell populations contribute distinct functional configurations that can nevertheless be represented within a common organisational framework, supporting a multicellular contribution to tissue-level functional organisation.

#### Functional organisation reveals continuous biological heterogeneity within conventional clinical classifications

To determine whether plaque functional organisation captures biologically meaningful disease organisation, we compared remodelling map-defined stage regions with conventional clinical classifications. Samples partitioned into Early, Intermediate, and Late map regions exhibited clear separation along the dominant axis of variation (Supplementary Figure 7). Plaque remodelling map-defined groups demonstrated clear ordering along PC1 and principal-curve pseudotime, whereas conventional disease labels showed greater overlap (Supplementary Figure 7). In contrast, PC2 and residual variation showed minimal association with organisation consistent with disease progression (Supplementary Figure 7). Next, we projected independent clinically annotated plaque patient datasets spanning asymptomatic, symptomatic and ruptured plaque into the remodelling map. Patient samples possessed coherent organisation (Figure 3A), with asymptomatic plaques mapping closer to early-stage healthy/control regions and ruptured plaques mapping closest to late-stage unstable regions (Figure 3A).

**Figure 3.**
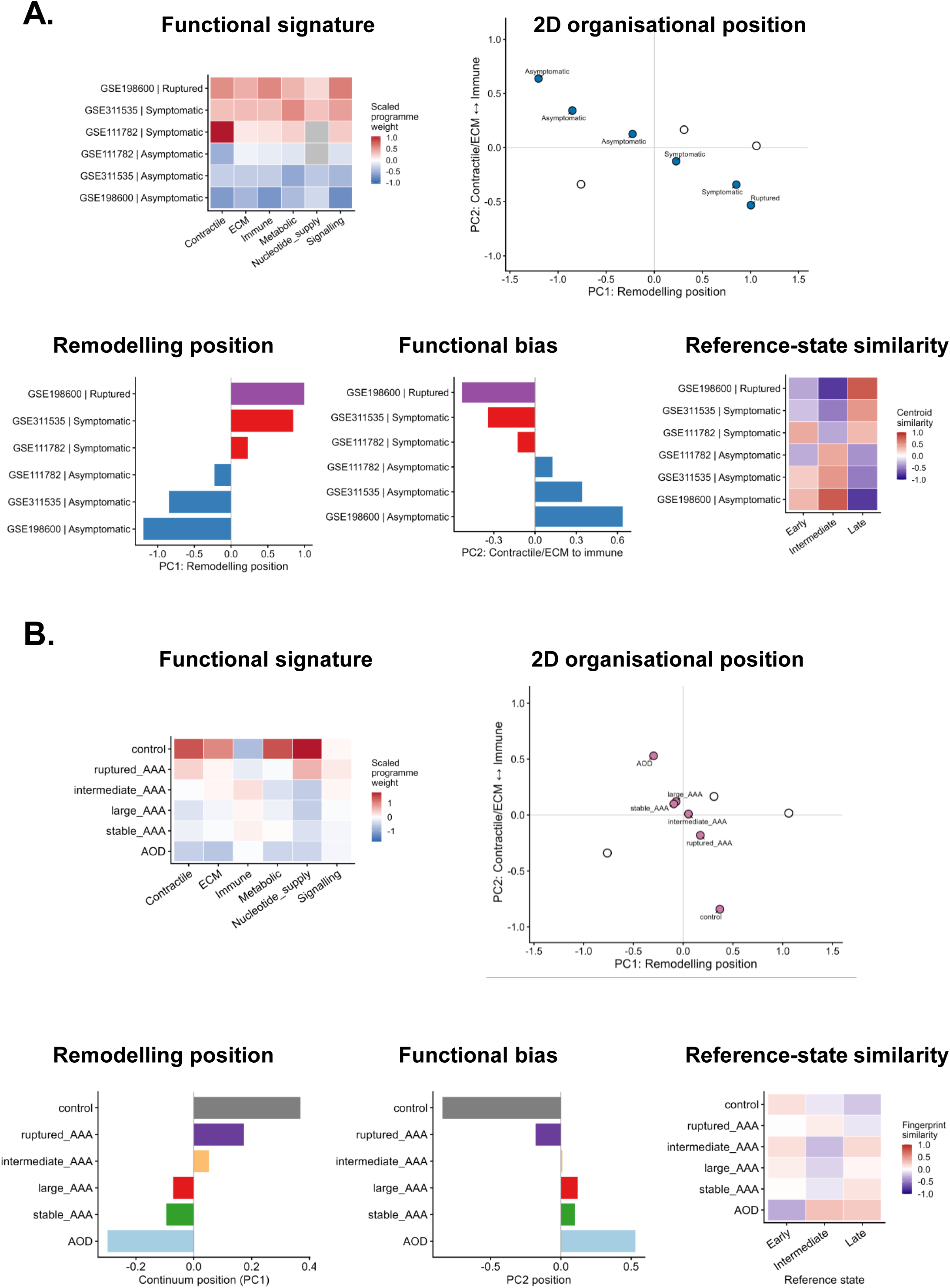
Functional organisation unifies clinical samples across atherosclerosis and AAA. **A)** Independent clinical plaque samples were organised within the reference plaque remodelling map. Functional signatures, 2D organisational position, remodelling position, functional bias and reference-state similarity revealed coherent differences across asymptomatic, symptomatic and ruptured plaques. **B)** Independent abdominal aortic aneurysm (AAA) transcriptomic datasets were projected into the same fixed reference plaque remodelling map. Functional signatures, 2D organisational position, remodelling position, functional bias and reference-state similarity revealed structured organisation across AAA samples and clinical subgroups.

We next determined whether this organisation was conserved across distinct vascular diseases. Independent abdominal aortic aneurysm (AAA) transcriptomic datasets occupied structured positions within the existing map (Figure 3B). Clinical aneurysm subgroups showed systematic relationships with the Early, Intermediate and Late continuum regions (Figure 3B).

### Genetic, pharmacological and dietary perturbations induce functional organisation reweighting

Independent genetic and pharmacological intervention datasets induced functional organisation rebalancing that resulted in positional movement within the shared plaque remodelling map (Figure 4A). Independent ApoE−/− datasets similarly demonstrated progressive movement towards advanced functional organisations following prolonged Western diet exposure (Figure 4B).

**Figure 4.**
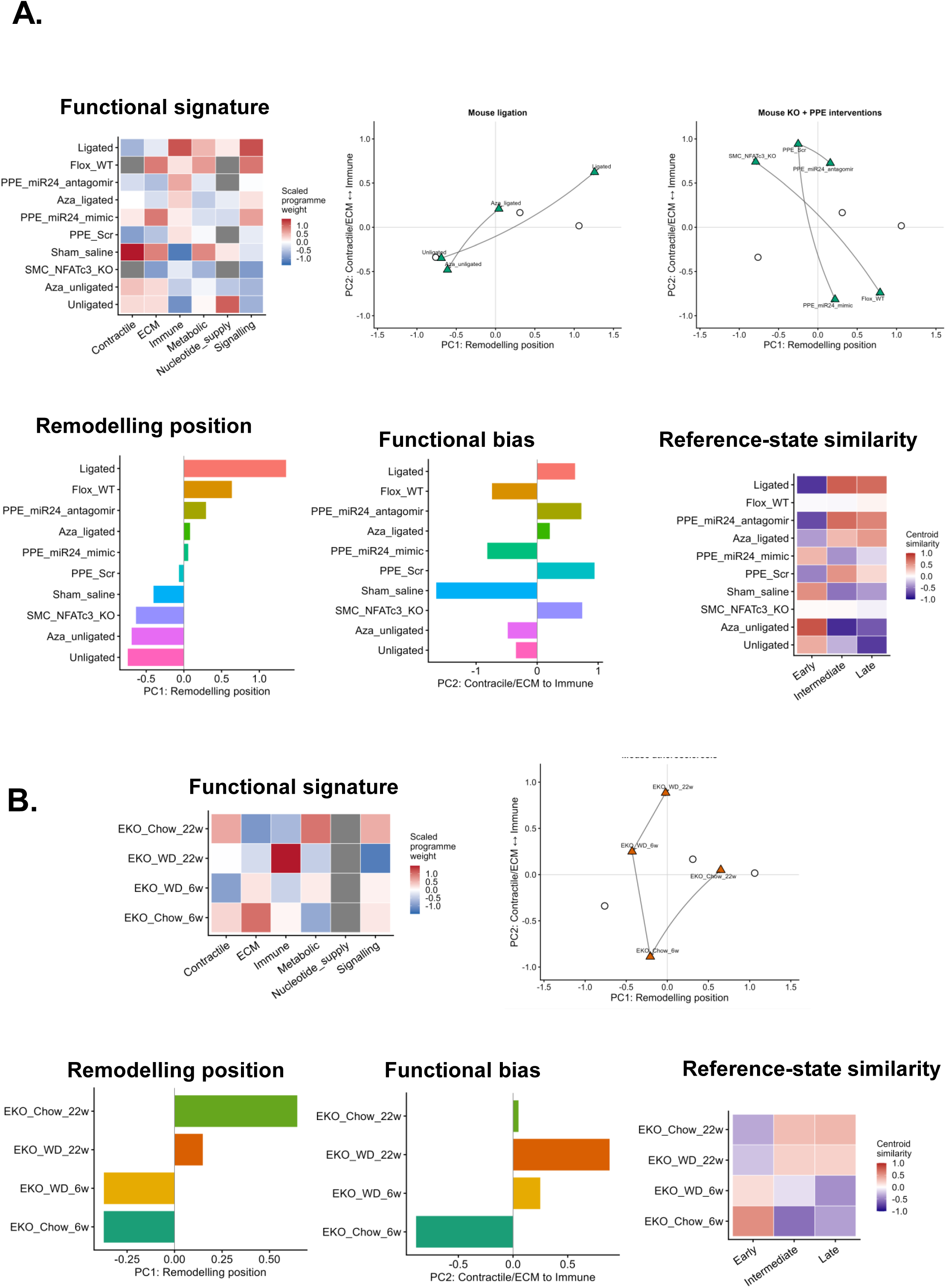
Experimental perturbation redirects vascular functional organisation. **A)** Independent genetic and pharmacological intervention datasets generated reproducible functional organisation rebalancing, quantified using functional signatures, 2D organisational position, remodelling position, functional bias and reference-state similarity. **B)** ApoE^-/-^ mice maintained on chow or Western diet for 6 or 22 weeks demonstrated progressive functional organisation rebalancing during disease progression. Increasing duration of Western diet exposure produced coordinated shifts in functional signatures, 2D organisational position, remodelling position, functional bias and reference-state similarity

## DISCUSSION

Cardiovascular diseases are commonly interpreted through molecular signatures, cellular phenotypes, and clinical classifications. While these approaches have yielded important mechanistic insights, they provide only a partial description of how complex multicellular tissues are organised during disease progression. It remains unclear whether a broader organisation exists that links these diverse observations across disease contexts. Our main finding is that vascular remodelling exhibits a reproducible and coordinated functional organisation that can be quantified from transcriptomic data. Despite substantial molecular complexity, much of the structure observed across vascular transcriptomic datasets was captured by the coordinated balance of a limited number of biological functions. The balance of these biological functions can be quantified as functional signatures that occupy positions in a shared remodelling map. Together, functional signature, remodelling position, functional bias, 2D organisational position and reference-state similarity provide complementary quantitative readouts of tissue-level functional organisation. Within this framework, disease progression, tissue heterogeneity, and therapeutic responses can be interpreted as functional organisation rebalancing and movement within this remodelling map.

There is a biological basis for such a shared vascular organisation. Thousands of genes contribute to vascular biology, but many participate in a relatively small number of coordinated biological functions. Because these functions are structured by the physical and functional requirements of vascular tissues, much of the observable transcriptomic variation may arise from changes in the relative balance of these functions. Such coordinated behaviour would be expected to generate a low-dimensional structure in which tissue position is determined primarily by the rebalancing of these biological functions.

The dominant functional organisation was preserved across pathway-retention thresholds, multiple within-cohort normalisation strategies, study-level exclusion, repeated sample subsampling and alternative dimensionality-reduction approaches, indicating that the observed structure is not attributable to a single analytical parameter or individual reference dataset. Independent datasets generated by different laboratories and platforms, and spanning different vascular territories, disease contexts and experimental perturbations, occupied coherent positions within the same reference map. Within this organisation, variation was defined primarily by the coordinated and nonlinear reweighting of structural, immune, signalling and metabolic functions, while a secondary organisational axis contrasted contractile/ECM organisation with immune activity. Importantly, functional organisation resolved biological heterogeneity that was not captured by conventional clinical classifications. Commonly used labels such as healthy, stable, unstable, and plaque capture only part of the underlying biological variation and corresponded to overlapping regions within the remodelling map.

Plaque-derived single-cell populations demonstrated coherent functional organisation. This supports the idea that tissue-level functional organisation reflects contributions from multiple cellular populations with distinct but coordinated functional states. This interpretation does not diminish the importance of molecular pathways or cellular phenotypes but provides a complementary and previously unrecognised organisational layer that emerges from collective cellular behaviour. This framework may also help explain the limited reproducibility of gene-level signatures frequently reported across vascular transcriptomic studies. Differences in disease stage, anatomical location, inflammatory burden, and cohort composition may generate distinct gene signatures while preserving a common underlying functional organisation. Functional organisation provides a transferable quantitative description of vascular tissue organisation that is less dependent on individual gene signatures.

Genetic, pharmacological and dietary interventions all induced structured repositioning within this organisation. Western diet-induced disease progression in ApoE⁻/⁻ mice followed functional organisation rebalancing and directional movement through the plaque remodelling map. These findings suggest that therapeutic responses may be represented as movement through the remodelling map. This raises the possibility that interventions could be evaluated by both the magnitude of transcriptional change and the direction of movement they induce within the remodelling map.

Although this framework was reproducible across diverse vascular contexts, its biological basis, clinical utility and potential to improve prediction remain to be established. Future studies should address three important questions. First, does functional organisation capture biologically meaningful information beyond existing molecular and cellular analyses? Second, can it provide a common quantitative framework for comparing tissues across molecular modalities and experimental systems? Third, can it improve prediction of plaque vulnerability, disease progression or therapeutic response?

Several limitations should be considered. First, the analyses are based predominantly on bulk transcriptomic datasets and therefore cannot fully resolve spatial organisation, cell-cell interactions, or local microenvironmental structure. Second, although functional organisation was highly reproducible across datasets and analytical approaches, the precise geometry of the organisational landscape remains influenced by dimensionality-reduction methodology. Importantly, functional organisation is not the product of dimensionality reduction. It is defined biologically as the coordinated balance of conserved biological functions. Dimensionality reduction provides one representation of this organisation, while functional signature, remodelling position, functional bias, 2D organisational position and reference-state similarity provide complementary quantitative readouts. Third, while the study identifies reproducible organisational structure and perturbation behaviour, it does not establish the molecular mechanisms responsible for functional organisation rebalancing. Fourth, although functional organisation rebalancing was associated with changes that were consistent with disease progression and intervention responses, causality remains to be established. Fifth, this study establishes the existence and reproducibility of functional organisation rather than its predictive performance. Whether functional organisation improves prediction of plaque vulnerability, myocardial infarction risk or therapeutic response will require prospective validation in appropriately phenotyped cohorts.

We propose that vascular disease can be understood at three complementary levels: molecular pathways, cellular phenotypes and tissue-level functional organisation. Our findings identify functional organisation as a previously unrecognised tissue-level property of vascular remodelling. The remodelling map provides a means of quantifying and comparing this property across diseases and experimental systems. Within this framework, disease progression, tissue heterogeneity and therapeutic response emerge as functional organisation rebalancing and directional movement through a shared remodelling map. Potentially, functional signatures provide a transferable representation that can be derived from transcriptomic, proteomic, imaging or spatial datasets. By providing a shared quantitative framework for describing multicellular tissue behaviour, functional organisation offers a foundation for integrating diverse molecular measurements across cardiovascular disease.

## Author contributions

Conceptualization: D.W.; Methodology: A.B., L.F., S.B., D.W.; Validation: A.B., L.F., D.W. Formal analysis: A.B., L.F., D.W.; Investigation: A.B., L.F., D.W.; Resources: D.W.; Data curation: D.W.; Writing-original draft: D.W.; Writing-review & editing: A.B., S.B., L.F., D.W.; Visualization: D.W.; Supervision: L.F., D.W.; Project administration: D.W.

## Sources of Funding

This work was funded by a UKRI Biotechnology and Biological Sciences Research Council Norwich Research Park Doctoral Training Partnership PhD Studentship awarded to A.B. (BB/T008717/1).

## Data availability

All datasets analysed in this study are publicly available through the Gene Expression Omnibus (GEO), with accession numbers provided in Table 1. Analysis code is available from the corresponding author upon reasonable request and will be deposited in a public repository upon publication.

## Disclosures

None

---

**Supplementary Figure 1.**
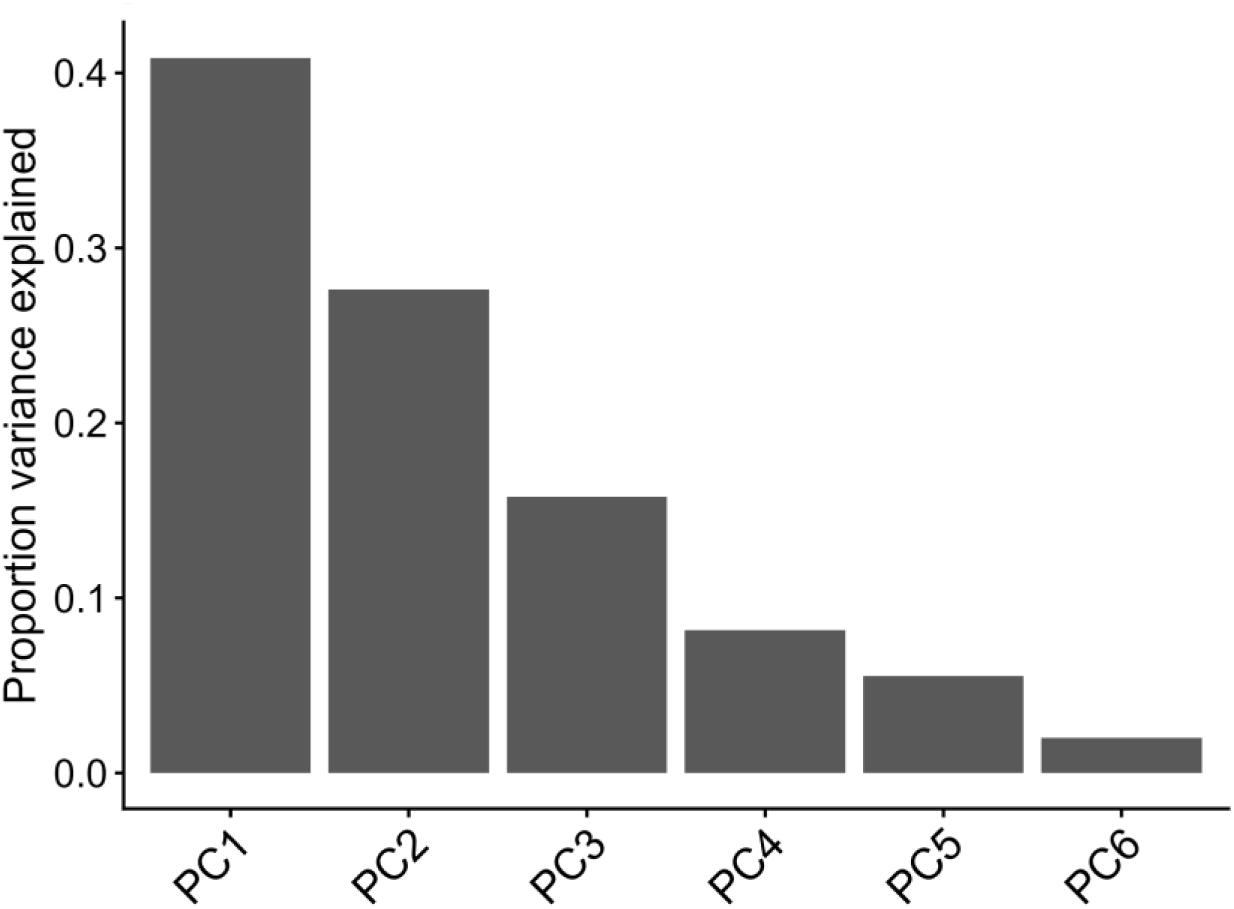
Variance explained by the reference plaque remodelling map. Principal component analysis of within-dataset z-scored biological-function weights demonstrating the variance explained by successive principal components. PC1 captures the dominant axis of functional organisation, with progressively smaller contributions from higher-order components. These data support representation of plaque remodelling within a low-dimensional functional organisational framework.

**Supplementary Figure 2.**
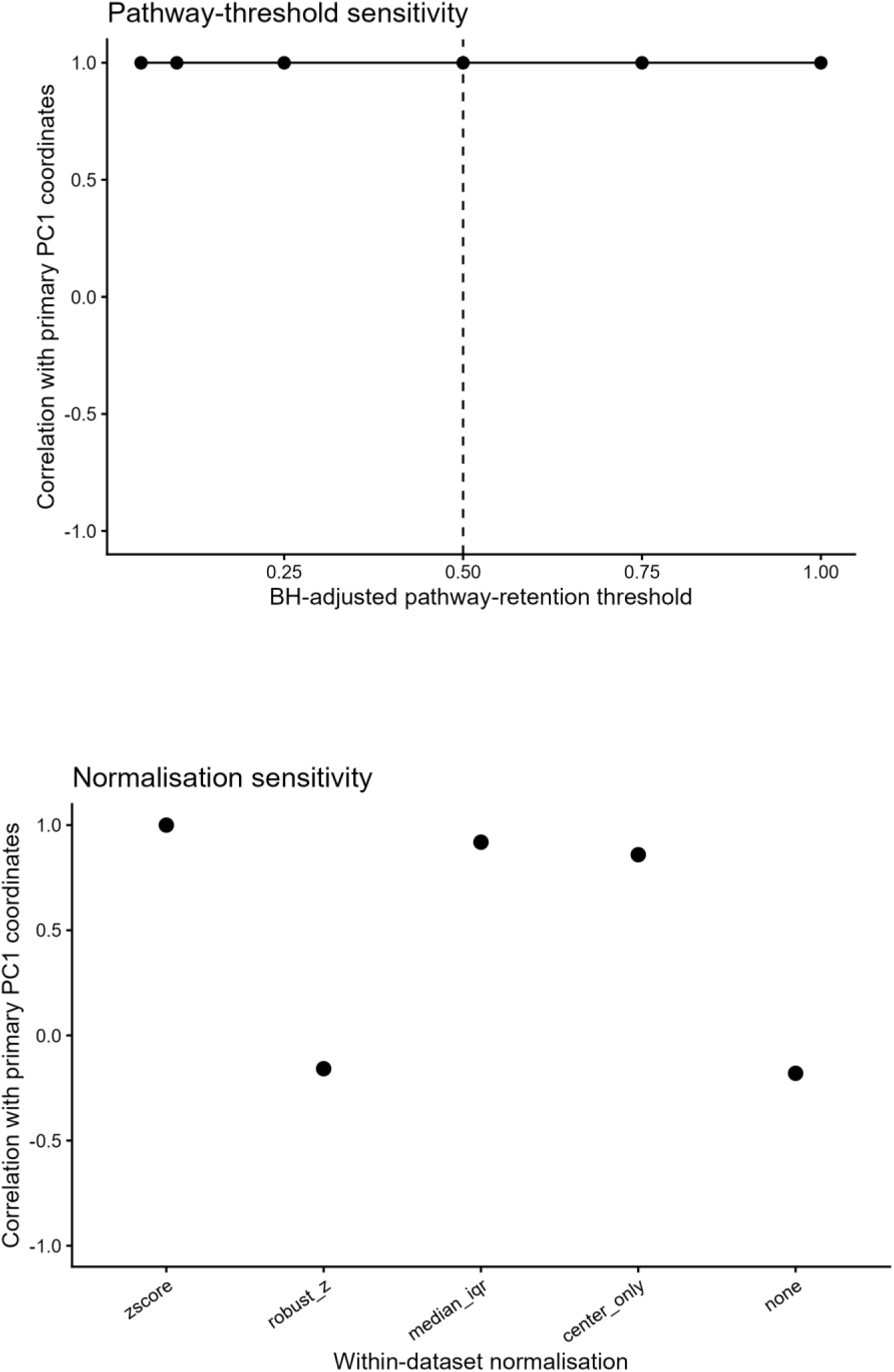
Sensitivity of the reference functional organisation to pathway-retention threshold and within-dataset normalisation. A) Pathway-threshold sensitivity analysis. The reference remodelling map was independently reconstructed across a range of BH-adjusted pathway-retention thresholds. Points show the correlation between PC1 sample coordinates obtained at each threshold and those from the primary analysis. The dashed line indicates the primary BH-adjusted pathway-retention threshold of 0.5. The dominant organisational axis was essentially unchanged across the range of thresholds examined. **B) Normalisation sensitivity analysis.** The reference remodelling map was independently reconstructed using alternative approaches for within-dataset normalisation, and PC1 sample coordinates were compared with those obtained using the primary within-dataset z-score transformation. Z-score normalisation, median–IQR scaling and centring alone strongly preserved the dominant organisational axis, whereas robust z-score transformation (median–MAD scaling) and omission of normalisation produced substantially weaker correspondence with the primary solution.

**Supplementary Figure 3.**
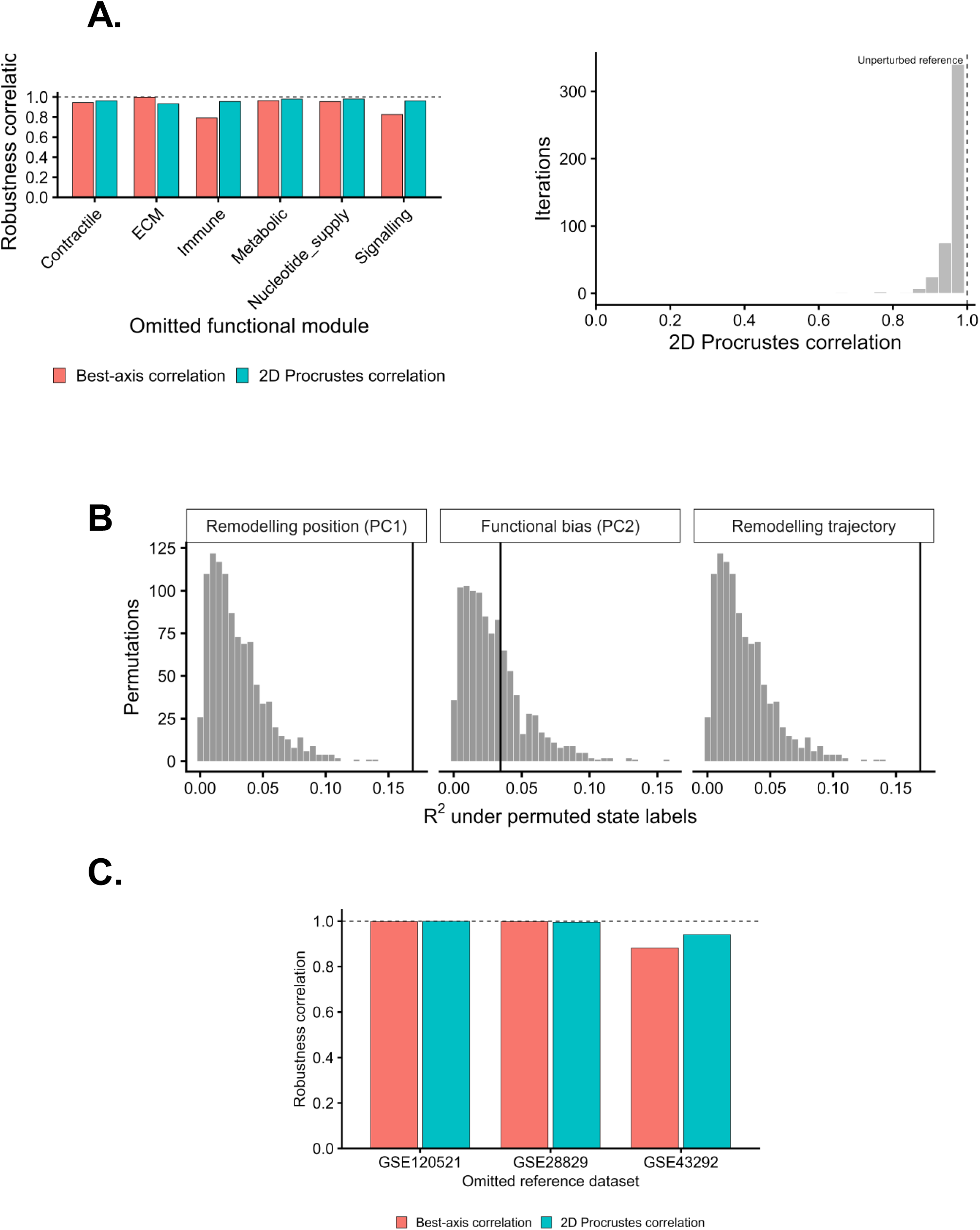
Functional organisation is robust to perturbation of functional composition, weighting and reference-dataset composition. **A)** Robustness of the reference functional organisation to perturbation of its biological-function representation. Left, each biological function was sequentially omitted and the reference map reconstructed. Bars show correspondence with the unperturbed reference quantified by best-axis correlation and two-dimensional Procrustes correlation. Right, the relative weighting of biological functions was randomly perturbed and the reference map reconstructed across repeated iterations. The distribution shows two-dimensional Procrustes correlations between perturbed and unperturbed solutions; the dashed line indicates the unperturbed reference. **B)** Permutation testing of the association between sample state and functional organisation. State labels were randomly permuted and the variance explained (R²) by state calculated for remodelling position (PC1), functional bias (PC2) and principal-curve remodelling trajectory. Histograms show the null distributions generated by permutation; vertical black lines indicate the observed values. The observed associations with remodelling position and trajectory substantially exceeded those expected under random state assignment, whereas functional bias showed a weaker state association. **C)** Leave-one-reference-dataset-out analysis. Each constituent reference dataset was sequentially omitted, the functional organisational map independently reconstructed, and correspondence with the complete reference quantified by best-axis correlation and two-dimensional Procrustes correlation. The dominant organisational structure was strongly preserved following exclusion of each dataset, indicating that the reference organisation is not dependent on an individual cohort.

**Supplementary Figure 4.**
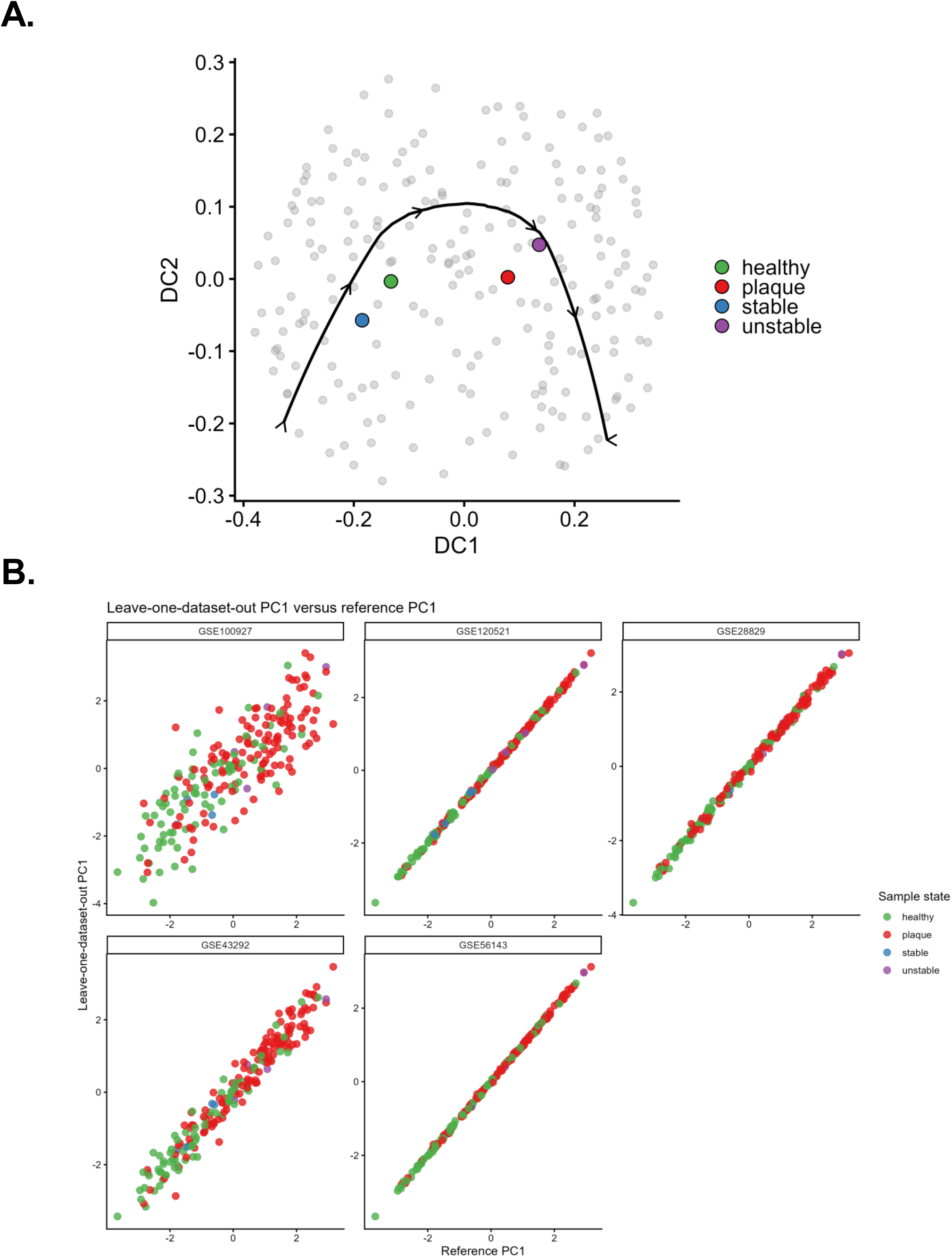
Functional organisation is preserved across analytical methods. **A)** Diffusion-map embedding of the reference datasets reproduces the ordered functional organisation identified by principal component analysis. Principal-curve fitting reveals the same dominant biological trajectory. **B)** Held-out/reference reconstruction analysis showing agreement of PC1 sample coordinates between leave-one-dataset-out reference maps and the complete reference, complementing the two-dimensional preservation analysis shown in Supplementary Figure 3C.

**Supplementary Figure 5.**
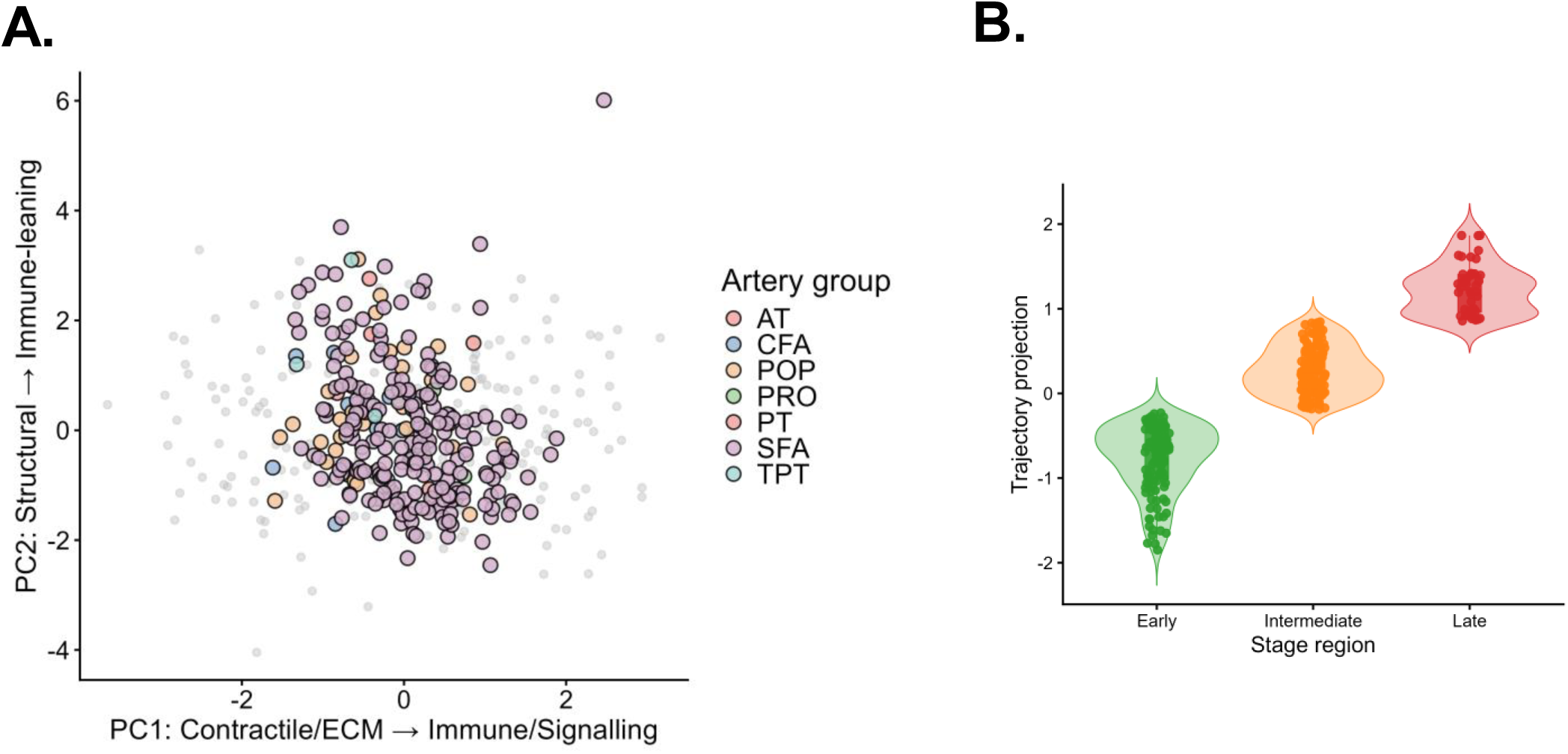
Functional organisation is conserved across vascular territories. **A)** Projection of independent peripheral artery transcriptomic samples onto the reference plaque remodelling map demonstrates preservation of the same functional organisation across multiple vascular territories. **B)** Distribution of peripheral artery samples along the reference continuum showing representation across Early, Intermediate and Late regions.

**Supplementary Figure 6.**
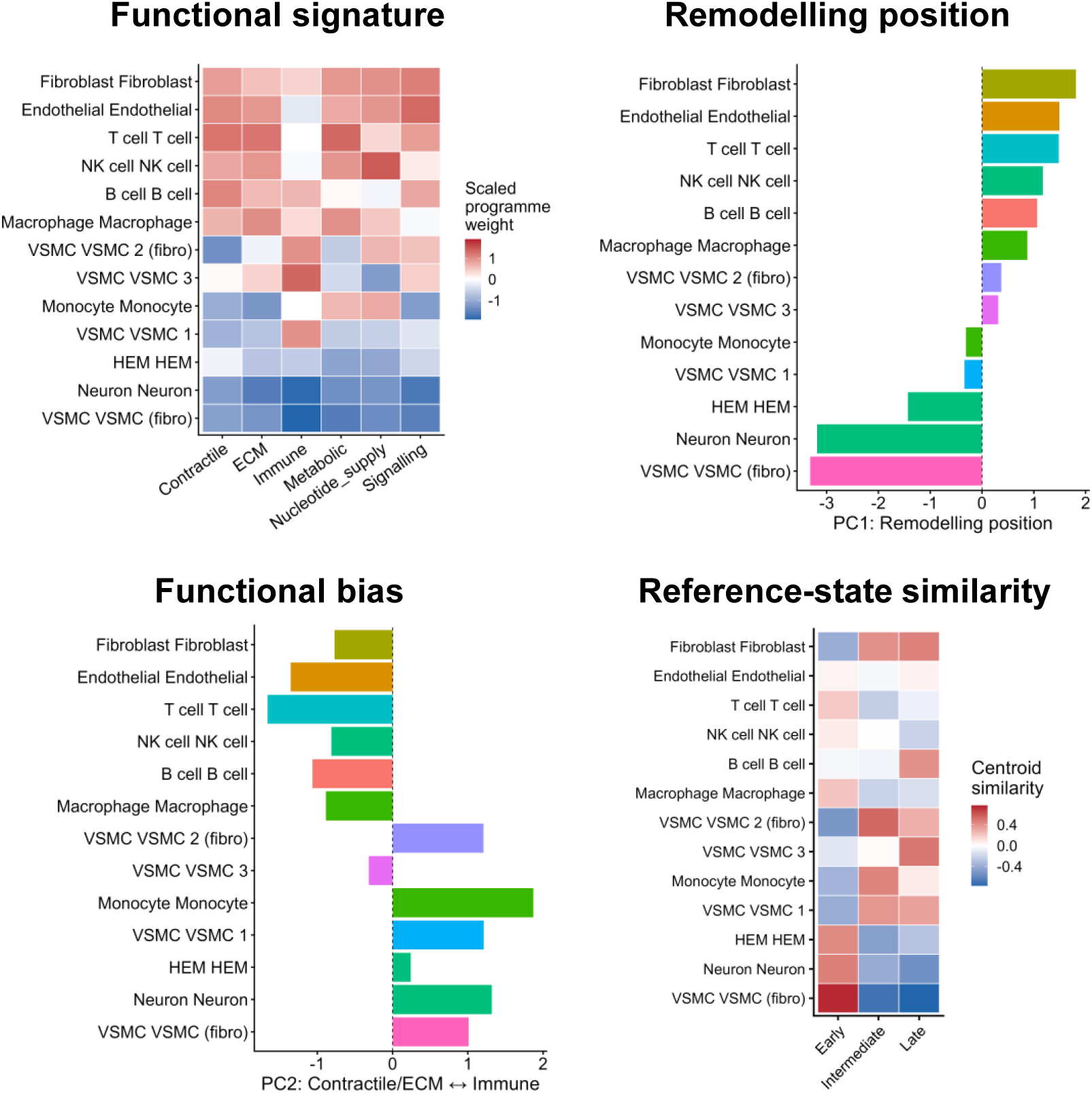
Functional organisation unifies cell populations. Functional signatures derived from independent single-cell RNA-sequencing differential expression profiles of plaque core and proximal cell populations demonstrate coherent functional signatures, remodelling position, functional bias and reference-state similarity.

**Supplementary Figure 7.**
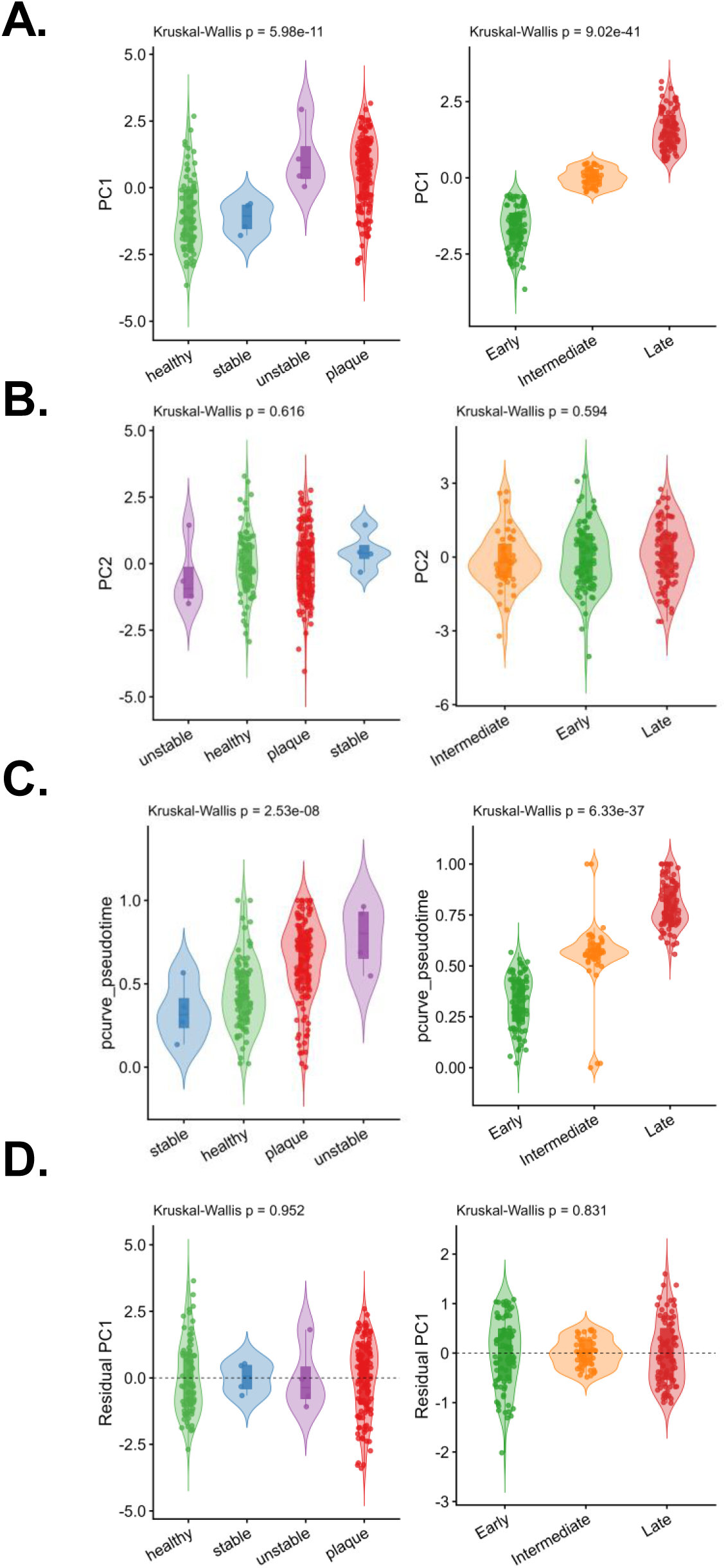
Disease progression is captured by the dominant axis of functional organisation. Independent clinical plaque samples were organised within the reference plaque remodelling map. Classification into Early, Intermediate and Late reference regions provided biologically coherent tissue-state stratification independent of conventional clinical labels. Violin plots comparing conventional clinical labels and reference-stage assignments. **A)** PC1 shows a significant association with conventional clinical labels and reference-stage regions. **B)** PC2 shows no significant association with either conventional clinical labels or reference-stage regions, indicating that orthogonal variation does not reflect disease progression. **C)** Principal-curve pseudotime demonstrates significant ordering across both clinical labels and reference-stage regions, consistent with progressive functional organisation rebalancing. **D)** Residual variation following removal of the dominant continuum shows no significant association with either clinical labels or reference-stage regions.

## Supplementary Methods

### Mathematical definition of functional organisation

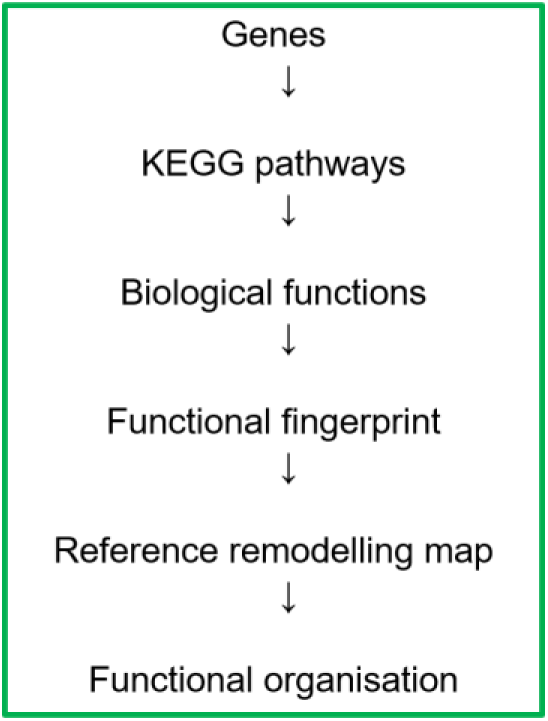

#### Conceptual definition of functional organisation

Functional organisation was defined as the coordinated relative balance of conserved biological functions across multicellular vascular tissues. Each sample was represented by six higher-order biological functions: Contractile, Extracellular Matrix (ECM), Immune, Metabolic, Nucleotide Supply and Signalling. These functions were derived from overlap-adjusted module-weighting scores calculated from KEGG pathway enrichment.

For sample *i*, the biological-function activity vector was defined as:

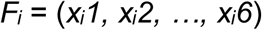

where *xᵢj* denotes the overlap-adjusted module-weighting score for biological function *j*. Together, these six measurements define the multidimensional functional state of each sample.

#### Functional signature

To enable comparison of relative functional organisation while reducing study-, platform- and scale-specific effects, biological-function scores were standardised independently within each source dataset or experimental cohort. For sample *i*, function *j* and dataset *d*:

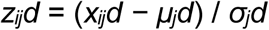

where *μⱼd* and *σⱼd* are the mean and standard deviation, respectively, of biological function j within dataset d. The resulting standardised vector

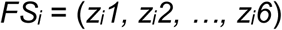

defines the functional signature of sample *i*. The functional signature therefore represents the relative balance of conserved biological functions within the sample rather than absolute transcriptional magnitude.

Where a biological function could not be estimated because insufficient pathway information was available, its biological value was retained as missing (NA). For PCA fitting or projection only, missing within-dataset standardised values were temporarily replaced by zero, corresponding to the within-dataset standardised mean. This computational substitution did not alter the stored biological-function scores.

#### Construction of the fixed reference remodelling map

The reference map was constructed exclusively from the human bulk atherosclerosis reference datasets. Let *Z_ref_* denote the analysis matrix of within-dataset standardised functional signatures used as input to PCA. With no additional centring or scaling applied during PCA, reference coordinates were calculated as

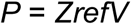

where *V* contains the PCA loading vectors and *P* contains the corresponding reference-sample scores. The loading matrix *V* was estimated once from the reference human atherosclerosis datasets and then fixed. No independent projection dataset contributed to estimation of the PCA basis.

#### Orientation of the reference axes

Because the signs of PCA components are mathematically arbitrary, component orientation was fixed after construction of the reference PCA. PC1 was oriented so that increasing values represented movement from healthy towards disease-associated reference organisation. PC1 therefore captures the dominant remodelling direction.

PC2 was oriented using only the biological-function loadings of the fixed human reference PCA. The sign was selected such that the Immune loading was greater than the mean loading of the Contractile and ECM functions. Increasing PC2 therefore represents increasing Immune weighting relative to mean Contractile/ECM weighting. PC2 is interpreted as a relative functional-balance axis rather than assigning biological meaning solely from the sign of an individual score.

#### Projection of independent datasets

Independent datasets were standardised within their own source dataset or experimental cohort and projected into the fixed human reference space without refitting the reference PCA. For an independent functional signature *FSnew*, its reference-map coordinates were calculated as:

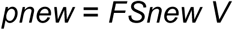

where pnew contains the projected principal-component coordinates, including PC1 and PC2.

Thus, independent datasets occupy a common coordinate system defined exclusively by the human atherosclerosis reference while retaining relative variation within their source cohort.

#### Remodelling position

Remodelling position was defined as the PC1 coordinate of a sample in the fixed reference PCA:

*RM_i_ = PC1_i_*

Because PC1 was oriented from healthy towards disease-associated reference organisation, increasing *RM* represents movement along the dominant direction of functional reweighting towards the disease-associated reference states.

#### Functional bias

Functional bias was defined as the PC2 coordinate:

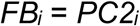

Increasing functional bias represents increasing Immune weighting relative to Contractile/ECM weighting within the orientation of the fixed reference PCA.

Two-dimensional organisational position

The joint PC1-PC2 coordinates retain both the dominant remodelling direction and orthogonal functional bias and define the two-dimensional organisational position:

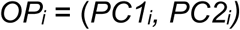

#### Principal-curve remodelling trajectory

A principal curve was fitted through the PC1-PC2 coordinates of the fixed reference samples to provide a complementary nonlinear description of the remodelling trajectory. Each reference or projected sample was associated with its position along the fitted curve, yielding principal-curve pseudotime (*PT*). Principal-curve pseudotime was used to characterise nonlinear progression through the reference organisation; it was not used as the primary definition of remodelling position, which was PC1.

For robustness analyses requiring direct comparison between independently reconstructed PCA solutions, rescaled PC1 (0–1) was used as the dominant remodelling coordinate rather than independently refitted principal curves.

#### Reference organisational states

Early, Intermediate and Late reference organisations were defined from the biological annotations of samples within the fixed human reference rather than from thresholds fitted to the projected datasets. For reference state (*s*), the centroid functional signature Cₛ was calculated as the mean standardised functional signature of samples assigned to that reference state:

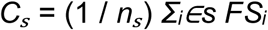

Reference-state signature similarity

Similarity between projected sample (*i*) and reference state (*s*) was quantified directly from their functional signatures using two complementary measures. Pearson correlation quantified signed pattern agreement:

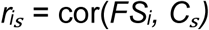

Cosine similarity quantified angular agreement between the two functional signatures:

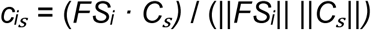

The two measures were combined with equal weighting:

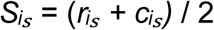

Combined similarities were converted to relative signature-match probabilities using a temperature-scaled softmax transformation:

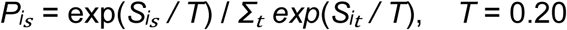

A fixed temperature parameter of *T* = 0.20 was used throughout to convert similarity scores into relative signature-match weights; it was not fitted to clinical outcomes.

These values quantify relative similarity of a sample’s functional signature to the Early, Intermediate and Late reference organisations. They are relative signature-match probabilities and are not calibrated probabilities of clinical state, disease outcome or future clinical events.

#### Quantitative representation of functional organisation

Functional organisation was represented using five complementary quantitative descriptors, which capture related but distinct aspects of the same organisational framework:

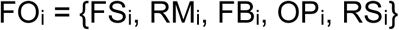

where FSᵢ is the functional signature; RMᵢ is remodelling position (PC1); FBᵢ is functional bias (PC2); OPᵢ is the two-dimensional organisational position (PC1, PC2); and RSᵢ denotes the vector of reference-state signature similarities or corresponding relative signature-match probabilities. These descriptors provide complementary quantitative readouts of the coordinated balance of biological functions within vascular tissue.

### Robustness analysis

#### Pathway-retention sensitivity

Dependence on the pathway-retention criterion was tested by reconstructing biological-function scores using BH-adjusted *P*-value thresholds of 0.05, 0.10, 0.25, 0.50, 0.75 and 1.00, while retaining the minimum requirement of three pathways per biological function. The reference PCA was independently reconstructed at each threshold and compared with the primary BH-adjusted *P* ≤ 0.50 solution after accounting for arbitrary PCA-axis orientation.

#### Normalisation sensitivity

Dependence on within-dataset normalisation was assessed by independently reconstructing the reference map using z-score standardisation (primary analysis), median-IQR scaling, median-MAD scaling, mean centring without scaling, or no within-dataset normalisation. Independently fitted solutions were aligned before comparison.

#### Leave-one-function-out analysis

Dependence on any single higher-order biological function was assessed by sequentially omitting each of the six functions and independently reconstructing the PCA from the remaining five. Because omission of a function can exchange or rotate the first two PCA components while preserving the underlying two-dimensional organisation, correspondence with the complete reference was assessed using best-matching-axis correlations and two-dimensional Procrustes alignment.

#### Random functional-weight perturbation

Dependence on the exact relative weighting of the six biological functions was assessed using 500 random perturbations. Multiplicative weights were drawn from a log-normal distribution with log-scale *SD* = 0.20 and re-centred within each iteration to a mean of one. This altered the relative contribution of the biological functions without systematically changing overall matrix magnitude. The PCA was reconstructed after each perturbation and correspondence with the unperturbed reference was quantified from sample-coordinate correlations, best-axis correspondence and two-dimensional Procrustes correlation.

This analysis tests dependence on the exact weighting of the higher-order biological functions; it does not randomly reassign individual KEGG pathways between functional dictionaries.

#### Leave-one-reference-dataset-out analysis

Each constituent human reference dataset was sequentially excluded and the reference PCA independently reconstructed from the remaining datasets. Correspondence with the complete reference was assessed using best-axis correlation and two-dimensional Procrustes correlation, thereby distinguishing preservation of the organisational subspace from arbitrary component swapping or rotation. Held-out samples could subsequently be projected into a reference space to which they had not contributed.

#### Stratified sample subsampling

Sensitivity to individual sample composition was assessed using 500 repeated subsampling iterations. In each iteration, 80% of samples were retained within each study-by-state stratum and the PCA was independently reconstructed. The resulting empirical distributions of correspondence metrics quantified sensitivity of the reference organisation to sample composition.

#### State-label permutation testing

To determine whether the relationship between functional organisation and clinical state exceeded that expected by chance, sample-state labels were randomly permuted 1,000 times while the measured functional signatures and PCA coordinates were retained. For each permutation, linear models quantified the proportion of variance (*R²*) in PC1, PC2 and the remodelling trajectory associated with the permuted state labels. The observed *R²* values were compared with the corresponding null distributions. Empirical permutation *P* values were calculated as:

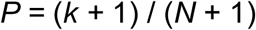

where k was the number of permuted *R²* values equal to or greater than the observed *R²* and *N* was the total number of permutations (1,000).

#### Alternative dimensionality-reduction analysis

To determine whether the dominant organisational structure depended specifically on PCA, an alternative diffusion-map representation was constructed from the same functional data. For the primary diffusion embedding, PCA was first performed on the six within-dataset standardised functional scores without additional centring or scaling, and the first five principal components were used as input to the diffusion analysis. The dominant diffusion coordinate was compared with the reference PC1 organisation. As an additional sensitivity analysis, diffusion embedding was performed directly on the six-function space without PCA preprocessing, and concordance between the resulting dominant diffusion coordinate, the PCA-pre-processed diffusion embedding and reference PC1 was assessed.

## Notes

### Competing Interest Statement

The authors have declared no competing interest.

